# Unveiling Mutation Effects on the Structural Dynamics of the Main Protease from SARS-CoV-2 with Hybrid Simulation Methods

**DOI:** 10.1101/2021.07.17.452787

**Authors:** P Gasparini, EA Philot, AJ Magro, JC Mattos, NESM Torres-Bonfim, A Kliousoff, RCN Quiroz, D Perahia, AL Scott

**Author notes:** Corresponding author: Ana Ligia Scott, Address: UFABC - Universidade Federal do ABC, Centro de Matemática, Computação e Cognição, Laboratório de Biofísica e Biologia Computacional, Campus Santo André, Avenida dos Estados 5001, Bairro Bangu, Santo André – SP, Brazil, CEP 09606-070. These authors contributed equally to this work (P Gasparini, EA Philot).

## Abstract

The main protease of SARS-CoV-2 (called M^pro^ or 3CL^pro^) is essential for processing polyproteins encoded by viral RNA. Macromolecules adopt several favored conformations in solution depending on their structure and shape, determining their dynamics and function. Integrated methods combining the lowest-frequency movements obtained by Normal Mode Analysis (NMA), and the faster movements from Molecular Dynamics (MD), and data from biophysical techniques, are necessary to establish the correlation between complex structural dynamics of macromolecules and their function. In this article, we used a hybrid simulation method to sample the conformational space to characterize the structural dynamics and global motions of WT SARS-CoV-2 M^pro^ and 48 mutants, including several mutations that appear in P.1, B.1.1.7, B.1.351, B.1.525 and B.1.429+B.1.427 variants. Integrated Hybrid methods combining NMA and MD have been useful to study the correlation between the complex structural dynamics of macromolecules and their functioning mechanisms. Here, we applied this hybrid approach to elucidate the effects of mutation in the structural dynamics of SARS-CoV-2 M^pro^, considering their flexibility, solvent accessible surface area analyses, global movements, and catalytic dyad distance. Furthermore, some mutants showed significant changes in their structural dynamics and conformation, which could lead to distinct functional properties.

**Highlights:**

- Single surface mutations lead to changes in M^pro^ structural dynamics.
- Mutants can be more stable than WT according to the structural dynamics properties.
- M^pro^mutants can present a distinct functionality in relation to the wild-type.
- Potential viral markers for more pathogenic or transmissible SARS-CoV-2 variants.

## 1. Introduction

The 2019 coronavirus disease (COVID-19) pandemic is the most severe health crisis in the past 100 years. As of June 07, 2021, approximately 173 million cases have been reported worldwide, with more than 3.72 million deaths (WHO Coronavirus (COVID-19) Dashboard). In addition to the zoonotic agents responsible for Severe Acute Respiratory Syndrome (SARS) and Middle East Respiratory Syndrome (MERS), the novel coronavirus, SARS-CoV-2, is a spherical-enveloped virus with a positive-sense single-stranded RNA (+ssRNA) which belongs to the broad *Coronaviridae* family and the *Nidovirales* order [1, 2]. The SARS-CoV-2 genome comprises about 30,000 nucleotides, including open reading frames (ORFs) responsible for encoding structural and non-structural proteins. ORF1a and ORF1b are translated into two overlapping polyproteins, pp1a and pp1ab, respectively [1]. These polyproteins are processed by the viral proteases M^pro^ (main protease) and PL^pro^ (papain-like protease) into 16 non-structural proteins (NSPs). Other ORFs undergo discontinuous transcription, and subgenomic mRNAs are translated into other non-structural, accessory, and structural molecules, such as envelope (E), spike (S), membrane (M) and nucleocapsid (N) proteins [3, 4].

SARS-CoV-2 M^pro^ is a dimer composed of three domains (the catalytic domains I and II and the C-terminal domain III), with the catalytic dyad formed by the residues His41 and Cys145 located in a cleft between domains I (residues 10-99) and II (residues 100-182) [5]. Dimerization is essential for SARS-CoV-2 M^pro^ enzymatic activity since the NH2-terminal residues (N-finger) of each monomer interact with the Glu166 residue of the other monomer, which effectively contributes to the correct arrangement of the S1 pocket of the substrate binding site. Thus, the C- and N-terminus of the monomers form the dimeric interface of the active protein and are less flexible in comparison to their higher mobility in the free monomers [5]. This general protein architecture is highly conserved in M^pro^s of diverse coronaviruses [6]; therefore, despite the extensive mutagenesis of these viruses in general, these key proteins are well conserved [7] and, therefore, are good targets for preventing virus replication and proliferation and reducing the risk of mutation-mediated drug resistance in new viral strains. Thus, the inhibition of M^pro^ enzymatic activity could be an interesting strategy for exploring new therapeutic approaches to treat COVID-19. Indeed, non-structural proteins have been reported as potential targets for the design and development of antiviral agents against SARS and MERS [8].

Recently, six worrisome SARS-CoV-2 variants, commonly associated with a potential escape of the immune response and a more transmissible and infectious capacity, were first detected in the UK (Alpha, B.1.1.7) [9], in South Africa (Beta, B.1.351) [10, 11], in Brazil/Japan (Gama, P.1) [12], in India (Delta, B.1.617+) [13], in USA/California (Epsilon, B.1.429+B.1.427) [14] and in UK/Nigeria (Eta, B.1.525) [15]. Interestingly, the genomic analysis of these variants listed in the GISAID database (https:gisaid.org) [16] highlights a set of frequent mutations such as K90R, P108S, K236R, L220F and R279C, in the SARS-CoV-2 M^pro^. Also, Amamuddy et al. (2020) [17] identified other non-synonymous mutations in all M^pro^ domains, including substitutions in solvent accessible residues and some in the N-finger region. Moreover, these last authors demonstrated the collective effects of various M^pro^ mutations in several SARS-CoV-2 isolates using different approaches and techniques, as geometry calculations, cavity compaction analyses, molecular dynamics simulations, anisotropic network model (ANM) calculations, and coarse-grained Monte Carlo simulations, among others.

From a protein dynamics perspective, it would be worth investigating the structural effects of these amino acid substitutions on the stability, activation, reactivity, and catalytic activity related to SARS-CoV-2 M^pro^. Hence, in this work, we analyzed the molecular characteristics as solvent accessible surface areas, the conformation of the catalytic dyad, and the flexibility of the regions involved in dimerization and substrate binding in wild-type and different SARS-CoV-2 M^pro^ mutants. A set of conformations of these molecules was generated using *in silico* methods based on all-atom molecular normal modes calculation. The results indicated that specific single mutations in SARS-CoV-2 M^pro^ could cause important changes in structural and dynamical characteristics regarding collective movements, the solvent accessible surface area (SASA) of the dimer interface, energetically accessible conformations, and arrangement of the catalytic dyad. These changes may influence the function and stability of SARS-CoV-2 M^pro^ present in important viral variants and could contribute to the emergence of harmful strains of SARS-CoV-2. This work analyzed the wild-type (WT) SARS-CoV-2 M^pro^ and several mutants using a hybrid approach *in silico* protocol to understand the possible influence of single mutations on this essential viral protein. The results shown here indicated that some of these mutations affect the SARS-CoV-2 M^pro^ structural dynamics and may potentially alter the functional properties of this macromolecule.

## 2. Material and methods

A Hybrid Method (VMOD, a module of the CHARMM program) that combines Normal Mode Analysis and low-temperature Molecular Dynamics simulations (described in the next section) was used to analyze the global motions and conformational changes for both WT and mutants of SARS-CoV-2 M^pro^. The characterization of the protein structural dynamics was done with the following analyses: a) flexibility with the Root Mean Square Fluctuation (RMSF); b) analysis of collective modes of wild-type and mutants (correspondence/correlation calculation); c) conformational sampling using hybrid methods; d) distance analysis between the two residues of the catalytic dyad; e) SASA analysis for different regions; f) analysis of the frequency of occurrence of mutants in the patient data deposited in the GISAID database.

### 2.1. SARS-CoV-2 M^pro^ mutants modeling

A total of 98 SARS-CoV-2 M^pro^ structures deposited in PDB until November 2020 were selected for initial analyses. Wild-type and apo crystal SARS-CoV-2 M^pro^ structures with 1.91 Å resolution (PDB ID: 7C2Y [DOI: 10.2210/pdb7C2Y/pdb]) were used to generate 48 mutants with single point mutations [17]using PyMOL software [The PyMOL Molecular Graphics System, Version 2.0 Schrödinger, LLC]. The most likely mutant rotamers were selected using the PyMOL Mutagenesis tool. The residue protonation states WT and mutants of SARS-CoV-2 M^pro^ at physiological pH (pH= 7.0) were assigned in the PDB2PQR server [18]. The protonated structures were submitted to the Solution Builder module of the CHARMM-GUI web server [19–21] to generate the input files for the molecular dynamics simulation used to analyze the Normal Modes [22–24]. We followed the protocol as outlined below:

● Selection of a dimeric wild-type and apo SARS-CoV-2 M^pro^ structure in PDB.

● Selection of SARS-CoV-2 M^pro^ mutants described by Amamuddy et al, 2020 [17].

● Building 48 mutants using Pymol and protonation with PDB2PQR server (see Figure 1). It is important to highlight that the start point of this study is the mutants reported by Amamuddy et al. (2020) [17].

**Figure 1:**
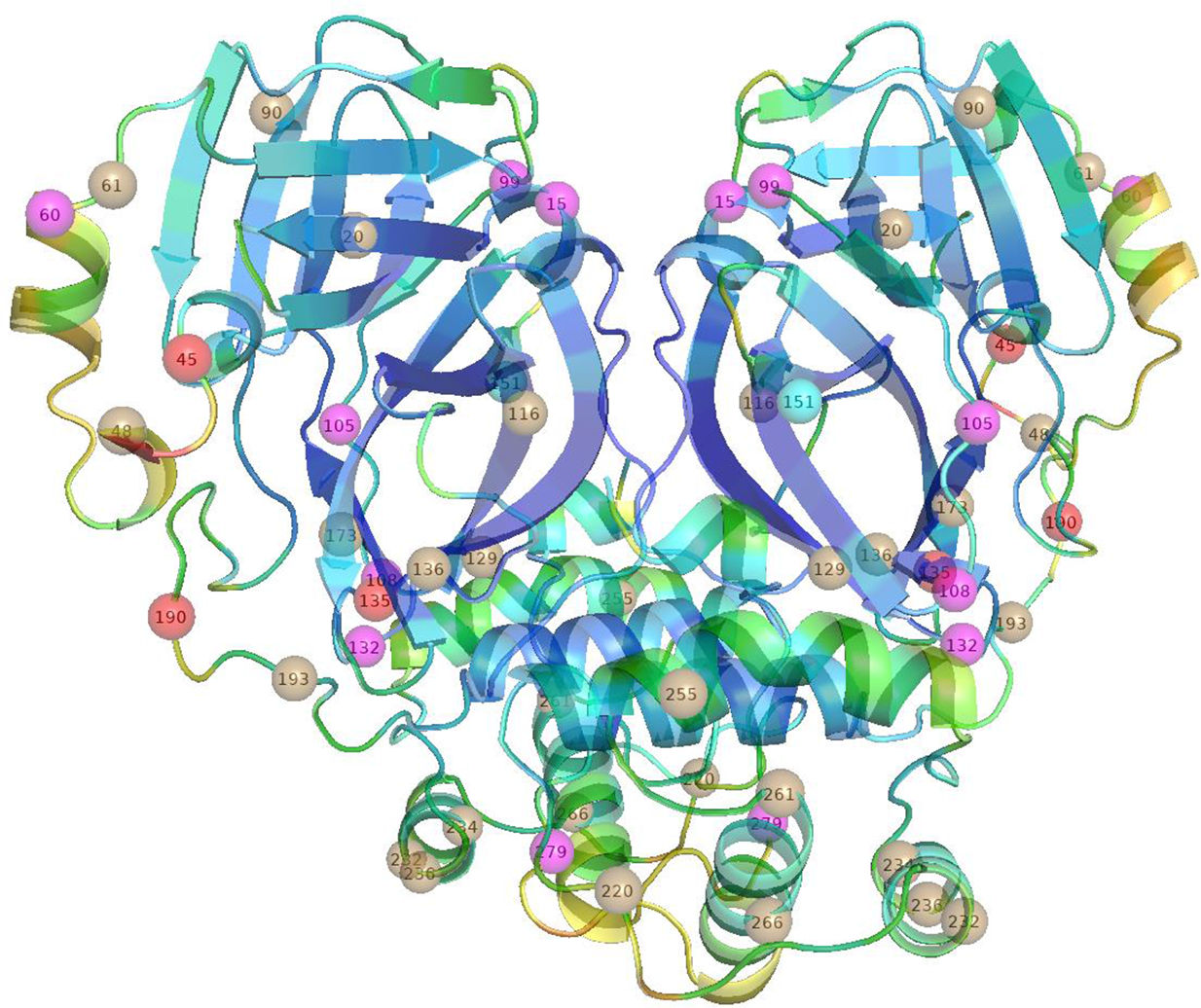
SARS-CoV-2 M^pro^cartoon representation with the mutated amino acid residues along the molecular tertiary structure. The backbone color corresponds to the RMSFs values for the WT where blue and red represent more rigid and flexible respectively. The positions of mutation for the mutants selected by RMSF (described in Table 2) are highlighted by spheres colored according to the characteristic of mutation: a) conservative (no charge change) in brown, b) charge inversion in cyan, c) polar to apolar without charge in red, d) special residue in magenta. Figure generated in PyMOL.

● Energy minimization of the WT and mutants of SARS-CoV-2 M^pro^.

● Calculation of the first 18 lowest frequency modes with an all-atom approach. The lowest frequency normal modes represent the collective motions of the protein.

● Analysis and selection of the mutants with a significant difference of RMSF compared to wild-type. We considered four important regions [5, 17] presented in Table 1 to accomplish this analysis.

**Table 1:** Important regions of the SARS-CoV-2 M^pro^ used as reference to compare the RMSFs and select the most significant mutants [5, 17].

| Region | Residues | Functionality |
| --- | --- | --- |
| **Catalytic Dyad (CatDy)** | His41, Cys145 | Responsible for substrate cleavage |
| **Substrate binding (Bsite)** | His41, Met49, Gly143, Ser144, His163, His164, Met165, Glu166, Leu167, Asp187, Arg188, Gln189, Thr190, Ala191, Gln192 | Region that undergoes conformational modifications to binding |
| **Dimerization (Dinterf)** | Arg4, Ser10, Gly11, Glu14, Asn28, Ser139, Phe140, Ser147, Glu290, Arg298 | Dimer Interface |
| **N-Finger (N-Fi)** | Ser1, Gly2, Phe3, Arg4, Lys5, Met6, Ala7, Phe8, Pro9 | Important for the Dimer Stability and Catalysis |

● Generation of energetically relaxed conformations along the lowest frequency normal modes by carrying out a low temperature MD and energy minimizations (the first 6 modes).

### 2.2. Normal Modes Analysis (NMA) and conformational selection of the most significant mutations

Initially, the protein structure was minimized in two stages. In the first one harmonic constraints were applied and progressively decreased (250 to 0 kcal mol^-1^Å^-2^) and for each constraint value 500 steps of gradient conjugate (CG) were applied. After the CG minimization the constraints were removed and 2x10^5^ steps of adopted basis NeWTon Raphson (ABNR) were used with the convergence criterion of 10^-6^ kcal mol^- 1^Å^-1^ RMS to energy gradient. The normal modes were calculated using the DIMB (Iterative Mixed-Basis Diagonalization) module [24] implemented in CHARMM software [25, 26] considering all the atoms of the protein and a force field corresponding to the CHARMM36m [27].

A set of the first six normal modes describing the internal movements were computed to ensure that all relevant large amplitude motions were included in our analysis. This is a sufficient set to ensure that all relevant large amplitude movements are included in our analysis. The modes corresponding to global rotation and translation were discarded. In the treatment of nonbonded interactions the Van der Waals (VDW) ones were computed up to 5 Å and a switching function was used to approximate these interactions until 9 Å. A distance-dependent dielectric constant (e=2ri,j) was employed to treat the shielding of electrostatic interactions by the solvent. In the work described here such a function has a good performance to predict realistic intrinsic global movements for all structures (WT and mutants of SARS-CoV-2 M^pro^).

### 2.3. Conformational space sampling along the collective modes

To obtain energetically allowed displaced structures along selected normal modes, we used the VMOD module from CHARMM [24] that makes use of harmonic restraints applied only to the Cα atoms. The restraints are set to target a given position along a specific normal mode vector. The MD and energy minimization were carried out for all atoms, enabling complete freedom of motion of the protein, including side chains. The structures were displaced from -1.0 to +1.0 Å (Mass Weighted Root Mean Square - MRMS values) along modes with steps of 0.2 Å, resulting in 11 intermediate structures for each mode. The displacements along the modes are achieved by using a series of low temperature MD simulations followed by energy minimizations. The procedure is similar to the one used by Batista et al. (2011) [28] and it is briefly described here. At each stage, the force constant of the harmonic restraining potential (Kd) was increased to ensure that the desired normal mode displacement was obtained. This procedure ensures that the structure slowly converges to the desired displacement. The restrain force Kd value was increased from 1,000 to 10,000 kcal·mol^-1^ Å^−2^ during successive 10 ps MD simulations. Velocities were assigned at random corresponding to a temperature of 30 K in the MD. The Berendsen thermostat with a coupling constant of 0.1 ps was used in all low temperature simulations.

Using a low-temperature dynamic allows for a more efficient conformational search than using only energy minimizations. A final MD simulation was achieved with a Kd value of 20,000 kcal·mol^-1^ Å^−2^ followed by 1,000 steepest descent and 1000 conjugate gradient steps of minimizations to reach the final displacement target along the mode.Using this procedure 66 structures were generated (11 structures per mode x 6 modes) for each molecular system considered.

### 2.4. Distance between the catalytic dyad atoms

According to Ramos-Guzmán et al. (2020) [29], the reaction mechanism proposed to proteolysis catalyzed by SARS-CoV-2 M^pro^ involves two steps, the acylation and deacylation steps. Briefly, in the first one the Sγ atom of Cys145 achieves nucleophilic attack to substrate peptide bond, before that occurs the formation of catalytic dyad ion pair by proton transfer from Sγ atom to the Nε atom of His41. Lastly, in the deacylation process, the bond between the M^Pro^ and the substrate is broken by hydrolysis that enables the enzyme to participate in a new catalytic cycle [29]. Regarding the first step of the reaction, the distance between the histidine nitrogen atom (Nε) and the cysteine sulfur (Sγ) was calculated to check its variability in the set of structures generated along the normal modes considered.

### 2.5. Solvent accessible surface area (SASA)

The CHARMM program was used to calculate the Solvent Accessible Surface Area (SASA) for the set of structures generated along the normal modes. We have reported the SASA values to the entire protein (SASAt - total), hydrophobic residues (SASAhp), dimeric interface residues (SASAdim), substrate binding site residues (SASAsb), His41 and Cys145 residue.

### 2.6. Mantel test

The lowest frequency normal modes calculated for WT and mutants of SARS-CoV-2 M^pro^ may not be completely redundant or possess matching. Several works in the literature show and report that a simple mutation can alter the motions present in the mutant, causing the emergence of new modes or the loss of some motion present in the WT. This same phenomenon can be noticed in the case of the SARS-CoV-2 M^pro^.

Comparisons between the movements of the WT and the chosen mutants of SARS-CoV-2 M^pro^ were performed using the protocol described in the Mantel test. In this protocol described by Louet et al. [29], motions in 3D space were analyzed by calculating 2D matrices, exchanging the relative displacements of all pairs of Cα atoms between two structures [30, 31]. Briefly, the method consisted of calculating 2D matrices reflecting the relative displacements of all pairs of Cα atoms between two structures. For each mode, these matrices were calculated from the two structures obtained at a displacement amplitude of 1.0 Å in each direction. The correlation coefficients between two matrices were then calculated with Mantel’s test [30]. With this method, it was assumed that two 2D maps sharing a correlation coefficient greater than 0.6 described highly related motions in cartesian space.

### 2.7 Statistical analysis

The RMSF arithmetic average was calculated for each analyzed region (N-finger, catalytic dyad, substrate binding and dimerization) of the WT and mutants of SARS-CoV-2 M^pro^. The selected mutants in which the average RMSF of at least one chain (A or B) exceeded the threshold of 10% compared to the RMSF values of the control group (WT).

The data obtained related to the potential energy, distance between the catalytic dyad atoms (Cys145, His41), and SASA from (i) binding site, (ii) hydrophobic residues, (iii) dimer interface residues, and (iv) total molecular surface were compared between the 48 mutants and WT of SARS-CoV-2 M^pro^ using ANOVA (Analysis of Variance) followed by Dunnett’s test (with significance at p ≤ 0.05). All statistical analyzes were performed using the multcomp package of ANOVA [32] implemented by R-Studio package [33]. The Heatmap, Violin and Boxplot representations were generated in Python using the Pandas [34, 35], NumPy [36], Matplotlib [37] and Seaborn [38] libraries.

### 2.8 Analysis of dimer interface

We used the PDBe PISA (Proteins, Interfaces, Structures and Assemblies) to investigate the interactions in the dimer interface for the WT and those mutants of SARS-CoV-2 M^pro^. Using the PISA web server, we calculate the solvation free energy gain upon formation of the interface, interface area and number of potential hydrogen bonds and salt bridge [39]. Complementary to the PISA results, we applied the TKSA-MC [40] to estimate the contribution of electrostatic interaction to free energy for the polar residues, especially those that are localized in the interface of the dimer (Arg4, Glu14, Glu240 and Arg298). The TKSA-MC calculates protein charge–charge interactions via the Tanford–Kirkwood Surface Accessibility model with the Monte Carlo method for sampling different protein protonation states. We used these two softwares to improve the discussion about the stability of the dimer for 4 mutants showing significant SASA and Potential Energy reduction (see Table 2).

**Table 2.**
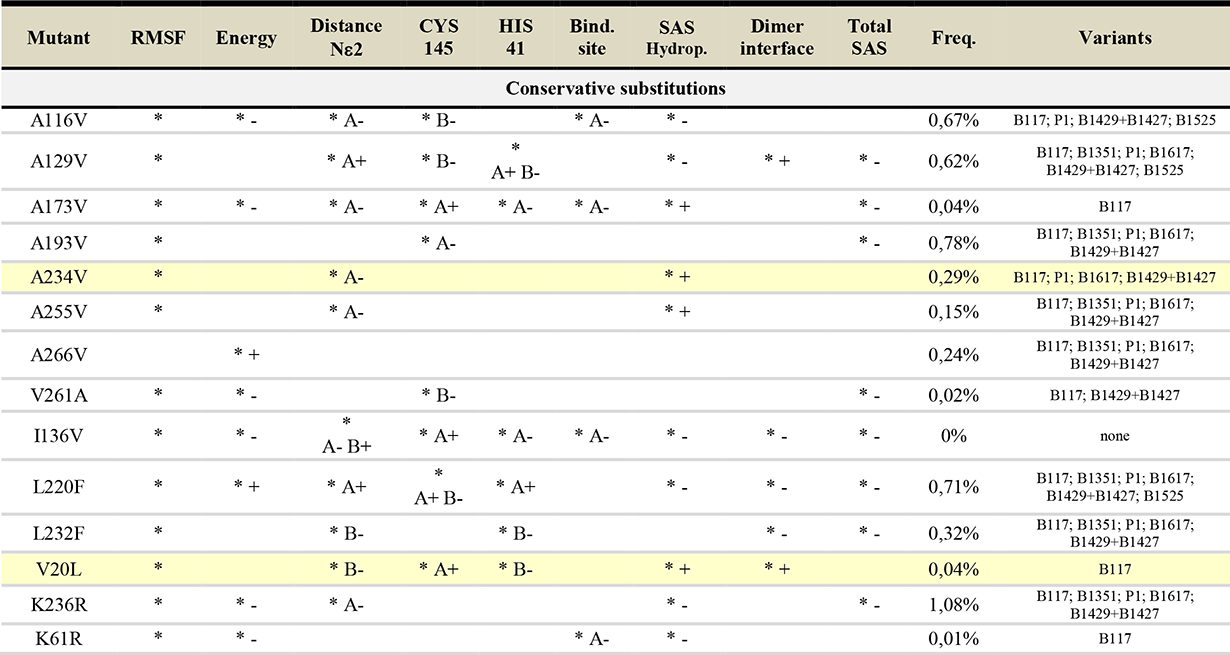
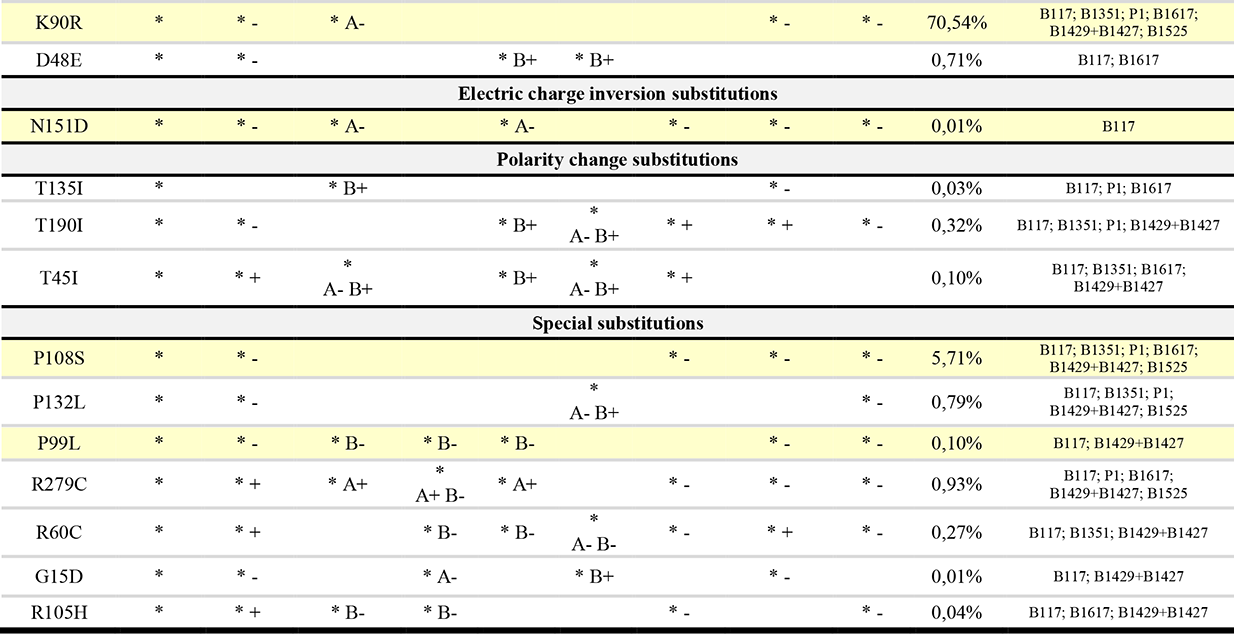
The table lists the mutants presenting at least three structural characteristics with significant variations compared with WT where the data are grouped by: 1) no charge change: apolar to apolar (AP -> AP: A->V, V->A, I->V, L->F, V->L), positive polar to positive polar (P(+) -> P(+): K->R, R->H), negative apolar to negative apolar (P(-) -> P(-): D->E); **2) charge inversion:** neutral to negative polar (NT -> P(-): N->D); **3)polar to apolar without charge:** polar to apolar (P -> AP: T->I); **4) special residue:** special to neutral (spe -> NT: P->S), special to apolar (spe -> AP: P->L), negative polar to special (P(-) ->spe: R->C), special to negative polar (spe -> P(-): G->D). **Columns means: Mutant:** mutation and residue; **Res.:** type of mutation (residues); **RMSF:** Root mean square fluctuations of chain A and B; **Energy:** energy calculated of each mutant; **Distance Nε:** distance between **Sγ** atom of Cys145 and **Nε** atom of His41; **Cys145 and His45:** catalytic dyad; **Bind. site:** SASA substrate binding site region; **Hidrop. SASA:** SASA of hydrophobic residues; **Total SASA:** accessible total area; **Freq.:** frequency of each mutant on variants (%), by sequences deposited on GISAID (last access: 31/05/2021). **Variant:** variants which the mutant are present in GISAID. The used symbols to denote significant statistics of parameter versus WT: “**\*”** (presence of significant statistics); “**-” (**decreased parameter), “**+”** (increased parameter), “**A-”** (decreased parameter on chain A), “**B-” (**decreased parameter on chain B), “**B+”** (increased parameter on chain B). In yellow are highlighted the probable stable mutant dimers.

## 3. Results and discussion

### 3.1. General characteristics of the SARS-CoV-2 M^pro^ mutants identified in GISAID database

Several works have reported variants of SARS-CoV-2 with mutations in different regions of M^pro^ and their effect on its functionality [5, 17]. Thereby, in this paper we intend to study the collective motions for the wild-type and different mutant SARS-CoV-2 M^pro^ molecules shed some light on the contribution of a group of known amino acid substitutions on the dynamics characteristics of this essential viral protein. For this, initially we calculated the normal modes for the WT SARS-CoV-2 M^pro^and 48 mutants of SARS-CoV-2 M^pro^ (set 1) using an all-atom approach. Following, we analyzed the structural dynamics of these molecules based on their Cα flexibility of key regions, such as the N-finger, dimeric interface, substrate binding site and His41- Cys145 catalytic dyad. As shown in Figure 2, both chains A and B from mutants of SARS-CoV-2 M^pro^ present significant fluctuations/differences in comparison to the wild-type molecule considering the average Cα RMSF (root mean square deviation) of the mentioned key regions. Additionally, there is a clear variation of the flexibility even between the chains A and B from the WT and mutants of SARS-CoV-2 M^pro^ (Figures S1, S2 and S3). In chain B, it is evident that the substrate binding region and the catalytic dyad of the mutants presented, on average, a higher flexibility in comparison to the wild-type molecule. This asymmetric/distinct behavior between the chains in homodimeric enzymes were also identified by molecular dynamics simulations involving the SARS-CoV-2 M^pro^ [17] and SARS-CoVM^pro^ [41] as well as NMA calculation for HIV-1 protease [42] and DPP-IV diabetes related protein [43]. Thus, this analysis indicated that some mutants may present characteristics that can influence their catalytic activity, substrate binding affinity and/or dimer stability, as also shown previously by Amamuddy et al. (2020) [17].

**Figure 2:**
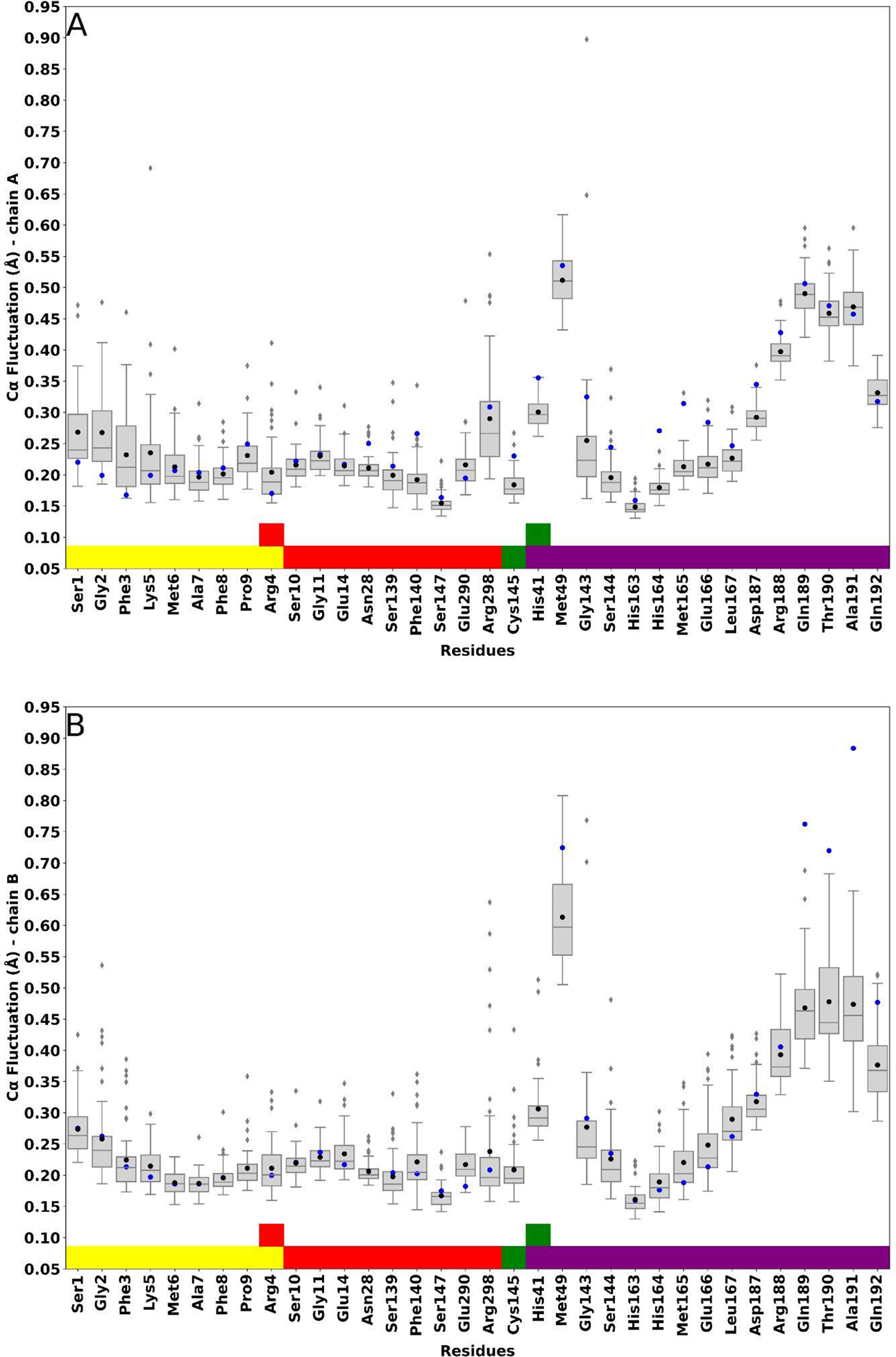
Boxplot RMSF of important regions of the SARS-CoV-2 M^pro^ reported in Table 1. (**A**) Chain A and (**B**) chain B. N-finger in yellow, dimeric interface residues in red, catalytic dyad in green and substrate binding site in purple. Black circle represents the mean of mutants and the blue circle represents the measure to WT. The residues Arg4 and Cys145 are not in ascending order in N-finger and catalytic dyad regions, respectively, to clarify that they are present in more than one region.

The initial analysis of the 48 SARS-CoV-2 M^pro^ mutants allowed the selection of 35 mutants with a high degree of flexibility (set 2), which were later evaluated considering some additional criteria as potential energy (Figure S4), SASAt (Figure S5), SASAhp (Figure S6A), SASAdim (Figure S6B), SASA of His41 and Cys 145 (Figure S7A and S7B), SASAsb (Figure S8), and catalytic dyad distance (Figure S9). Thus, this new screening of the calculated structures allowed to underscore 26 mutants of SARS-CoV-2 M^pro^ (set 3) with significant statistical variation in at least three of the mentioned criteria in comparison to WT SARS-CoV-2 M^pro^. Additionally, these 26 mutants from set 3 were divided into four groups according to the type of amino acid substitution, i.e., involving conservative substitution (when a given amino acid residue is replaced by a different one with similar biochemical properties), electric charge or polarity modification, and special cases in which occurs the inclusion/exclusion of amino acid residues with unique properties (cysteine, glycine, histidine or proline) (Table 2).

The most number of mutants from set 3 (15 structures) was classified in the first case, involving conservative replacement. In this group, besides significant Cα RMSF alterations, nine mutants showed relevant modifications of potential energy. More specifically, seven of these 15 mutants (D48E, K90R, I136V, A173V, L220F, and L232F) presented a negative variation of the potential energy and only one (L220F) showed a positive variation of the potential energy in relation to the WT SARS-CoV-2 M^pro^. From these structures, the amino acid substitution D48E was the unique which did not present a significant negative variation in at least one of the considered additional criteria (SASAdim, SASAsb, SASAhp, and SASAt). Considering the occurrence of potential energy and SASAdim variation and at least one of the parameters related to surface exposition (SASAhp and SASAt), it is possible to point out five mutants of SARS-CoV-2 M^pro^ with statistically significant differences in comparison to the WT SARS-CoV-2 M^pro^ (K90R, I136V, A173V, L220F, and L232F). Particularly, the mutation K90R is present in the variants B.1.1.7 (first detected in the UK), B.1.351 (first detected in South Africa), P.1 (first detected in Brazil/Japan), B1.617 (first detected in India), B.1.429+B.1.427 (first detected in USA/California), and B.1.525 (first detected in UK/Nigeria) and also is markedly prevalent in the SARS-CoV-2 M^pro^ sequences deposited in the GISAID database (around 70% of frequency). Another relatively prevalent mutation identified in the GISAID database was the K236R amino acid substitution, which presents 1,08 % of frequency and could be found in the variants B.1.1.7, B.1.351, P.1, B1.617, and B.1.429+B.1.427. In this case, there is no significant difference of the SASAdim in comparison to the wild-type molecule, however it was verified SASAt and SASAhp alterations. Curiously, the mutation I136V is the only one which presents significant variation in the SASAdim, SASAt and SASAhp, but was found in none of the variants from the GISAID database.

The other SARS-CoV-2 M^pro^ amino acid substitutions from set 3 (11) were grouped according to the electric charge/polarity alteration or associated to the inclusion/exclusion of cysteine, glycine, histidine, or proline amino acid residues. Only one substitution lead to the introduction of a negative-charged amino acid residue (N151D); it is important to highlight that this mut SARS-CoV-2 M^pro^ molecule was just found in the variant B.1.1.7 and showed a significant variation of the potential energy, SASAdim, SASAt and SASAhp in relation to the WT SARS-CoV-2 M^pro^. Another three mutants of SARS-CoV-2 M^pro^ presented polar (not electrically charged) to non-polar amino acid substitutions involving the change of threonines by isoleucine residues (T45I, T135I, and T190I). In this case, the amino acid substitution T190I was found in four variants (B.1.1.7, B.1.351, P.1, and B.1429+B.1427) and could be considered that with the more important alterations compared to WT SARS-CoV-2 M^pro^, since it introduces important modifications related to the potential energy, SASAdim, SASAsb, SASAhp, and SASAt. Finally, it were identified in the GISAID database seven amino acid substitutions related to the inclusion/exclusion of cysteine, glycine, histidine, or proline amino acid residues (G15D, R60C, P99L, R105H, P108S, P132L, and R279C). Here, it is possible to underline the amino acid substitution P108S, which can be identified in six variants (B.1.1.7, B.1.351, P.1, B1.617, B.1.429+B.1.427, and B.1525) and is also the most prevalent among the seven mutants of this subgroup, with 5,71 % of frequency considering the SARS-CoV-2 M^pro^ sequences deposited in the GISAID database. In addition, the substitution P108S promotes significant alterations of the SASAdim, SASAsb, and SASAt, as well as the substitutions R60C and R279C. However, the substitutions R60C and R279C are associated with an increase of the potential energy in comparison to the WT SARS-CoV-2 M^pro^, whereas the substitution P108S decreases the potential energy.

### 3.2. Modifications of the SARS-CoV-2 M^pro^ dimeric interface induced by amino acid substitutions at protein surface

Up to now, the solved crystallographic structures of M^pro^ molecules from coronavirus indicate they are structurally very similar and basically dimeric. The M^pro^ protomers from coronavirus consists of three well-characterized domains, being two with a chymotrypsin fold and a third extra helix domain, which is essential for dimerization of this viral protease [44–48]. Also, since the mid 2000s, a series of scientific works has confirmed that the dimerization process is fundamental to the full catalytic activity of the main proteases from coronavirus [5]. Hence, it is reasonable to suppose that the combination of low levels of potential energy, lesser SASA values and larger dimeric interfaces could favor the protein stabilization and, in the specific case of the mutants of SARS-CoV-2 M^pro^, to affect the activity of these enzymes. About this issue, Hu et al. (2009) [49] showed that two specific and neighbouring mutations at the SARS-CoV-1 M^pro^ dimeric interface lead this molecule to assume distinct oligomeric conformations. In fact, this work concluded that certain key amino acid residues control the SARS-CoV-1 M^pro^ dimerization and also suggested the dimeric stability of this protease is heavily dependent on the extent and integrity of the contacts between the monomers. In general, the calculated mutant structures from set 3 showed an expected tendency of SASAt decreasing associated with a negative variation of the potential energy. Although the mutants of SARS-CoV-2 M^pro^ do not present amino acid substitutions at their dimeric interfaces, some of the calculated mutant structures showed wider contact areas in relation to the wild-type molecules. More specifically, four surface amino acid substitutions from set 3 (K90R, P99L, P108S, and N151D) (Table 2) allowed to reach the mentioned parameter combination (lower energy, smaller SASAt, larger dimeric interfaces) considering their respective calculated structures. Thus, these mutants of SARS-CoV-2 M^pro^ were carefully examined to assess their differences to the WT SARS-CoV-2 M^pro^, particularly those related to the dimeric interfaces and the influence of the K90R, P99L, P108S, and N151D amino acid substitutions in the conformational space accessed by the mutant molecules.

The PDBePISA web server indicated that some calculated K90R and N151D mutant structures presented a higher dimeric interface contribution to the total solvation free energy (Δ^i^G) not including the effect of satisfied hydrogen bonds and salt bridges across the interface) in relation to the wild-type molecule and the other mutants (Figure 3A). Therefore, at least these two mutants are able to assume oligomeric conformations which are kept by a dimeric interface more hydrophobic than the remaining wild-type and mutant structures generated by the computed normal mode displacements. On the other hand, as shown in Figure 3A, there is a visible concentration of wild-type molecules which reach Δ^i^G values around -11.0 kcal/mol whereas the calculated mut SARS-CoV-2 M^pro^ structures showed mostly higher Δ^i^G values. Hence, the K90R, P99L, P108S, and N151D mutants can also form, with a higher probability, dimeric interfaces which offer a lesser hydrophobic contribution to the Δ^i^G in comparison to the WT SARS-CoV-2 M^pro^. Indeed, the Figure 3C indicates the K90R, P99L, P108S, and N151D mutants can potentially establish at their dimeric interfaces a higher number of hydrogen bonds than the wild-type protease, indicating thus the formation of more stable and specific contacts between the protomers. An example regarding this difference between the wild-type and mutant calculated structures is the influence of the mutation P99L. In this case, it was possible to generate a dimer presenting approximately ten hydrogen bonds at the interface more than the wild-type calculated structure with the highest number of hydrogen bonds connecting its monomers. Moreover, according to the TKSA-MC server [40], the electrostatic contribution to the total free energy (ΔGqq) of four amino acid residues present at the mut SARS-CoV-2 M^pro^ dimeric interfaces (residues) considering both protease chains was clearly distinct. As shown in Figure S10 (Supplementary Material), the ΔGqq values related to the mutant chains A were lower in comparison to the corresponding chains of the calculated wild-type structures. Otherwise, this difference is not observed for the chains B from the calculated wild-type and mutant K90R, P99L, P108S, and N151D structures. Although the ΔGqq dimeric interface electrostatic contribution is based on the amino acid residues Arg4, Glu14, Glu290 and Arg298, this finding corroborates the results calculated by the PDBePISA web server that also indicated the prevalence of polar dimeric interface interactions in several mutant structures. Indeed, Ding et al. (2005) [50] also highlighted that hydrophobic contacts and electrostatic interactions play major roles in the binding of a SARS-CoVM^pro^ dimerization inhibitor based on affinity capillary electrophoresis experiments. Then, based on the preceding findings, it is feasible to suggest that the surface amino acid substitutions K90R, P99L, P108S, and N151D were able to induce a higher density of structures with different dimeric interfaces (and consequently distinct oligomers), which also showed larger extensions and a considerable number of polar contacts. Thus, the comparison with the calculated wild-type structures indicates that the mutations in question may lead to more stable SARS-CoV-2 M^pro^s.

**Figure 3:**
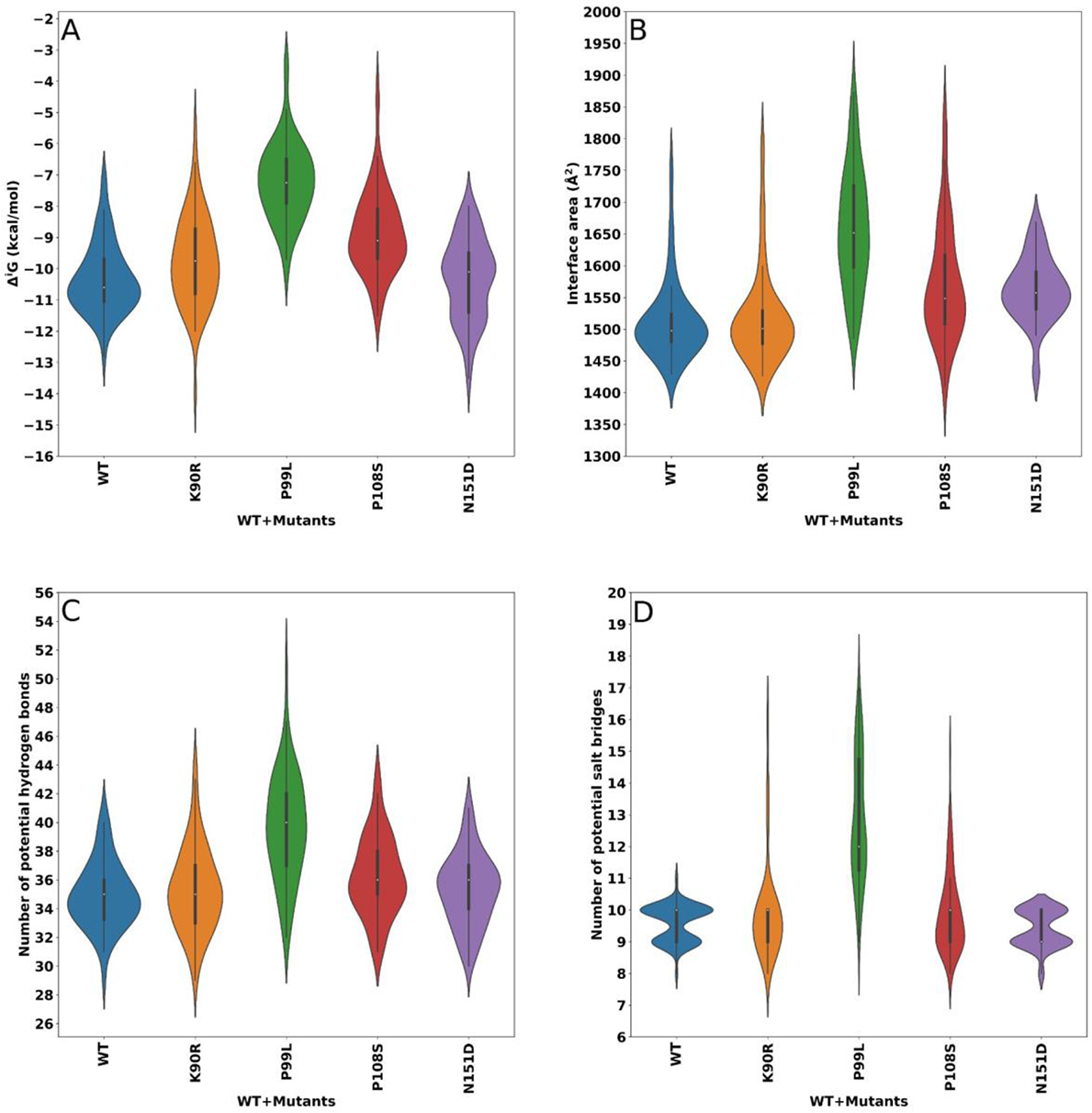
Dimeric interface analysis using PISA web server. (**A**) Δ^i^G indicates the solvation free energy gain upon formation of the interface (kcal/mol). (**B**) Interface area (Å^2^). (**C**) Number of potential hydrogen bonds. (**D**) Number of potential salt bridges.

### 3.3. Amino acid substitutions X catalytic dyad distance

It is well known that some amino acid residues are essential to the dimerization and activity of the main proteases from SARS-CoV-1. Bacha et al. (2004) [51] were the first authors to demonstrate this fact by identifying a vital cluster of conserved serine residues at the neighborhoods of the SARS-CoV-1 M^pro^ active site. The substitution of these serine residues by alanines impaired significantly the enzymatic activity of this protease. Simultaneously to the previous work, Chou et al. (2004) [52] reported the importance of a salt bridge interaction between Arg4 and Glu290 for SARS-CoV-1 M^pro^ subunit association; under high salt concentration and low pH conditions this ionic interaction is missed leading to loss of dimerization and reduced activity of this CoV-1 protease. Hsu et al. (2005) [45] reported that the truncation of the residues 1 to 4 located at the N-terminal region promoted SARS-CoV-1 M^pro^ monomerization and substantially diminished its catalytic activity. Chen et al. (2008) [53] identified critical amino acid residues related to SARS-CoV-1 M^pro^ dimerization based on mutagenesis experiments. In this work, the authors found the Ser10 and Glu14 residues located at SARS-CoV-1 M^pro^ domain I are crucial to the formation of interface interactions, being highly conserved in several coronavirus main proteases.

Additionally, Chen et al. (2008) [54] reported that the amino acid substitution G11A led to a total monomer separation after crystallization of this mutant SARS-CoV-1. Next, Shi et al. (2008) [55] showed that the SARS-CoV-1 M^pro^ R298A mutation produced crystals with monomeric molecules. In this study, the authors highlighted that Arg298 is a key amino acid residue involved in the dimerization of this protease. Remarkably, Zhong et al. (2008) [56] demonstrated that the truncated-N-finger M^pro^ C- terminal domain from SARS-CoV-1 can remain in a monomeric form and also dimerize forming a new interface. However, in this case, the new dimer is inactive, thus confirming the N-finger relevance for the formation of catalytically active SARS-CoV-1 M^pro^ molecules. Also, Hu et al. (2009) [49] identified two close mutations (S139A and F140A) at the SARS-CoV-1 M^pro^ dimer interface which resulted in different conformations of the crystal structure of this enzyme.

Most of the previous works based their conclusions on amino acid residues close to the dimeric interface or directly involved in the formation of dimeric contacts of the SARS-CoV-1 M^pro^. However, our results indicated that the distinct conformational behaviour of the calculated K90R, P99L, P108S, and N151D mutants and wild-type SARS-CoV-2 M^pro^ structures are exclusively due to surface mutations since there was no alteration of the amino acid residues which compose the dimeric interface of these molecules. This is an interesting point, mainly considering the notable sequence and structural similarity between the SARS-CoV-1 and SARS-CoV-2 M^pro^ molecules. Indeed, it is possible to suppose that the surface K90R, P99L, P108S, and N151D amino acid substitutions may not only alter the dimeric interface but also the functionality of the SARS-CoV-2 M^pro^. From this perspective, it would be noteworthy to investigate how the structural alterations related to these mutations could affect the nucleophilic attack of the Sγ atom of the Cys145 to the substrate scissile peptide bond.

According to Ramos-Guzmán (2020) [29], multiscale simulation methods showed the most probable between the catalytic dyad amino acid residues (Sγ atom from and Nε atom from H41) is around 3.3 Å, considering a conformer which keeps the shortest possible distance of the C45 side chain to a bound substrate molecule. The analysis of different apo-form dimeric SARS-CoV-1 and SARS-CoV-2 M^pro^ molecules available at the Protein Data Bank (PDB) also showed similar catalytic dyad distances (3.5 to 4.3 Å) (Table S1). As shown in Figure 4, some of the calculated K90R, P99L, P108S, and N151D mutant structures were able to reach catalytic dyad distances below 4.0 Å. Therefore, these results indicated the proposed protocol involving displacement of the initial wild-type and mutant structures generated molecules with a catalytic dyad geometry prone to trigger the substrate catalysis.

**Figure 4:**
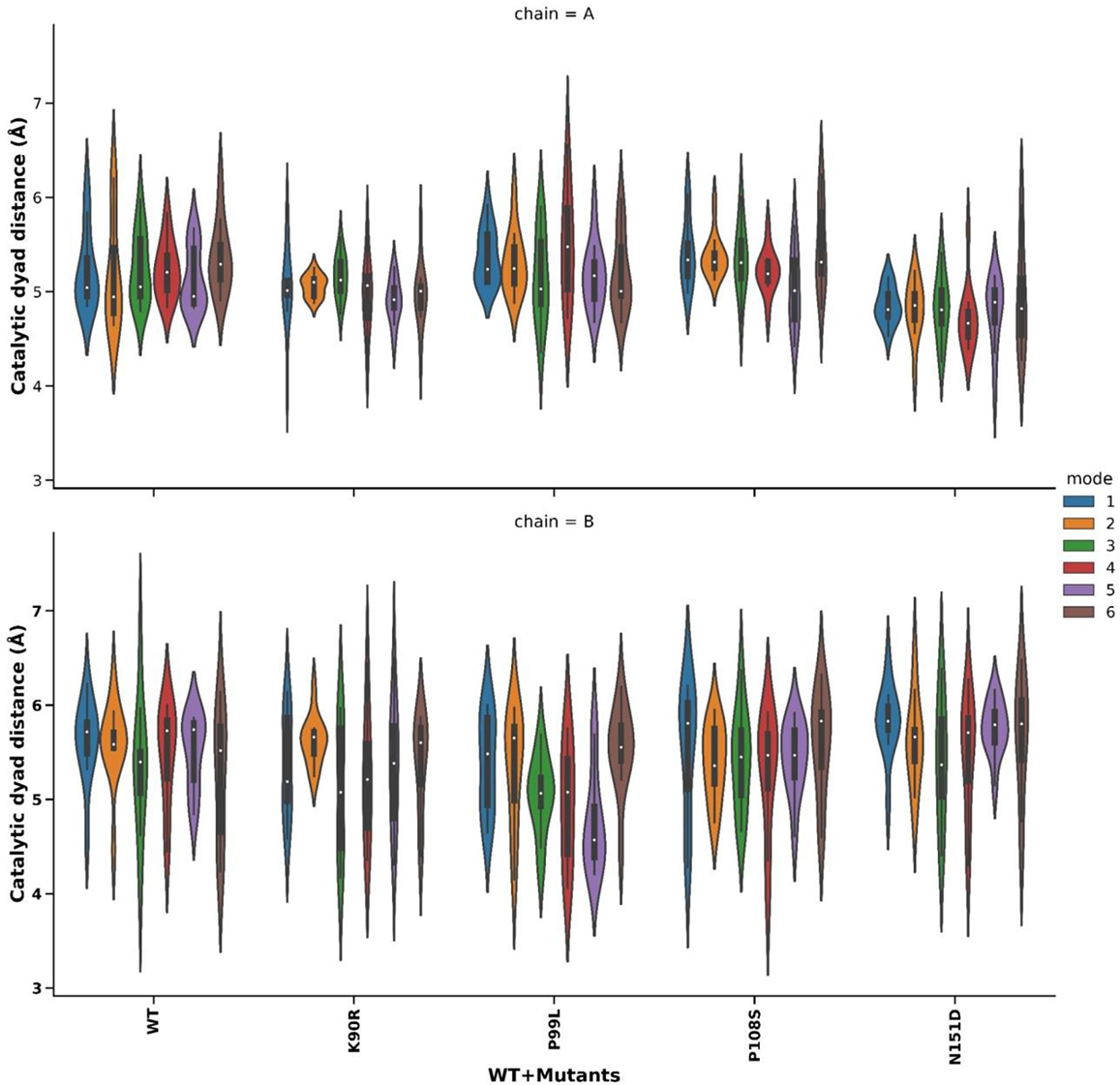
Catalytic Distance: Distance Violin plot by chain. Distance between the Cys 145 Sγ atom and His 41 Nε atom for both chains of the structures generated by VMOD. Chain A in blue and chain B in orange. The WT SARS-CoV-2 M^pro^ measures are highlighted by a red rectangle.

Remarkably, new layers of information emerge based on the analysis of the collective motions of the calculated wild-type and K90R, P99L, P108S, and N151D mutant structures. Only the modes 3, 4, and 5 from mutant and wild-type molecules present some conservation degree (Table S2). This finding is very significant since it demonstrates the four surface mutations in question modify the SARS-CoV-2 M^pro^ dynamics, a fact that is a strong indication of functional diversity. Besides, there is also a higher catalytic dyad distance variation in the protomers B from the mentioned four mutants. Except for the normal mode 6, all of the other normal modes (modes 1 to 5) allowed the generation of mutant protomers B with shorter catalytic dyad distances in comparison to the corresponding wild-type protomers (Figure 4). It is also noteworthy to highlight that the K90R, P99L, P108S, and N151D mutants presented movements which brought their protomer B catalytic dyads to distances close to theoretical and experimental values.

These results may be analyzed under the light of the work executed by Chen et al. (2006) [41], which employed MD simulations and mutational studies to demonstrate that only one protomer of the dimeric SARS-CoV-1 M^pro^ is catalytically active. Hence, the number of calculated mutant structures (K90R, P99L, P108S, and N151D) with catalytic dyads presenting closer distances to that adequate for nucleophile activation (the C145 Sγ atom) also indicated that one of the SARS-CoV-2 M^pro^ protomers (in this case, the protomer B) has a higher probability to reach conformations more appropriate for catalysis. Further, other important information comes from a more detailed comparison between the wild-type protease and the K90R, P99L, P108S, and N151D mutants. As verified in Figure 4, the modes which generate protomers B with catalytic dyad distances around or below 4.0 Å are the following: (i) 1, 2, 3, 4, and 6 for WT SARS-CoV-2 M^pro^; (ii) 1, 3, 4, 5 and 6 for K90R mut SARS-CoV-2 M^pro^, (ii) 2, 3, 4, 5, and 6 for P99L SARS-CoV-2 M^pro^, (iii) 1, 4 and 6 for P108S SARS-CoV-2 M^pro^; and (iv) 3, 4 and 6 for N151D SARS-CoV-2 M^pro^. However, as plainly indicated by the distribution of the calculated structures (Figure 4), the modes do not present the same probability of generating protomer B molecules with catalytic dyad distances adequate for catalysis reaction. As example, it is possible to compare the modes 5 from the protomers B from wild-type and P99L mutant molecules; in this case, the wild-type movement do not generate structures with catalytic distances close to 4.0 Å, whereas the movements correspondent to the mode 5 lead to a higher number of mutant conformers with catalytic dyad distances in the appropriate range for nucleophile formation. In truth, most of the collective motions related to the conformations with shorter catalytic dyad distances correspond to a set of different mixed bending/twisting movements, including bending motions between monomers and twist torsions around the axes of the monomers (Figure S11). Thus, the results shown here point to some hints that could help to identify eventual differences between the action mechanism of wild-type and mutant SARS-CoV-2 M^pro^ molecules. As highlighted by Goyal and Goyal (2020) several works have been based on the design of substrate binding pocket-ligands to inhibit the SARS-CoV-1 M^pro^, but none of them has reached clinical trials to date. Thus, a good therapeutic alternative to fight against SARS-CoV-2 could be to target the dimerization of its main protease. In this work, the importance of this process for SARS-CoV-2 M^pro^ is strengthened by the evident influence of surface mutations on the oligomeric conformation of this essential viral protease and, probably, to its molecular evolution and adaptation. Finally, the results shown here may also help to identify factors related to the emergence of more pathogenic or transmissible viral types.

## 5. Conclusions

This work compared the WT and mutants of SARS-CoV-2 M^pro^ using a normal modes-based protocol involving the analysis of simple parameters as Cα flexibility, potential energy, hydrophobic and electrostatic contributions to free energy, solvent accessible surface areas, and catalytic dyad distance. These parameters allowed the selection of potential structurally stable dimers, which were posteriorly analyzed and remarkably showed that single surface amino acid substitutions are able to induce significant dimeric interface changes. Also, the mutants showed a low conservation of collective motions in relation to the wild-type molecule according to the Mantel test. Additionally, the K90R, P99L, P108S, and N151D mutants presented modes with different probabilities of generating conformers with catalytic dyad prone to trigger the catalysis reaction in comparison to the wild-type protease. Therefore, this finding allowed us to suppose the mutants could present a distinct functionality in relation to the original SARS-CoV-2 M^pro^.

Finally, it is important to emphasize again the importance of the dimerization process for SARS-CoV-2 M^pro^ and the evident influence of surface mutations on the oligomeric conformation of this protein. Thus, this work may shed some light on the molecular evolution of this viral protease and the adaptation process of the SARS-CoV- 2 to their human hosts and help to identify markers related to more dangerous variants and strains of this virus.

## Appendix A. Supplementary data

Supplementary data to this article can be found online.

---

**Figure S1:**
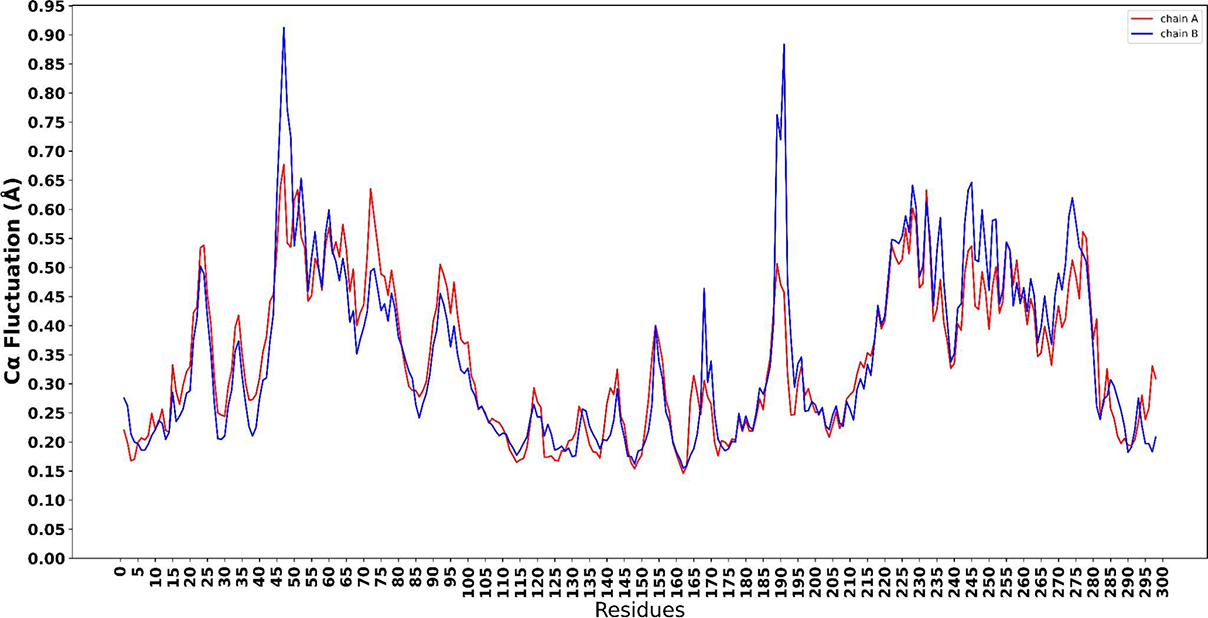
WT SARS-CoV-2 M^pro^ Cα fluctuation obtained by NMA. Chain A in red and chain B in blue.

**Figure S2:**
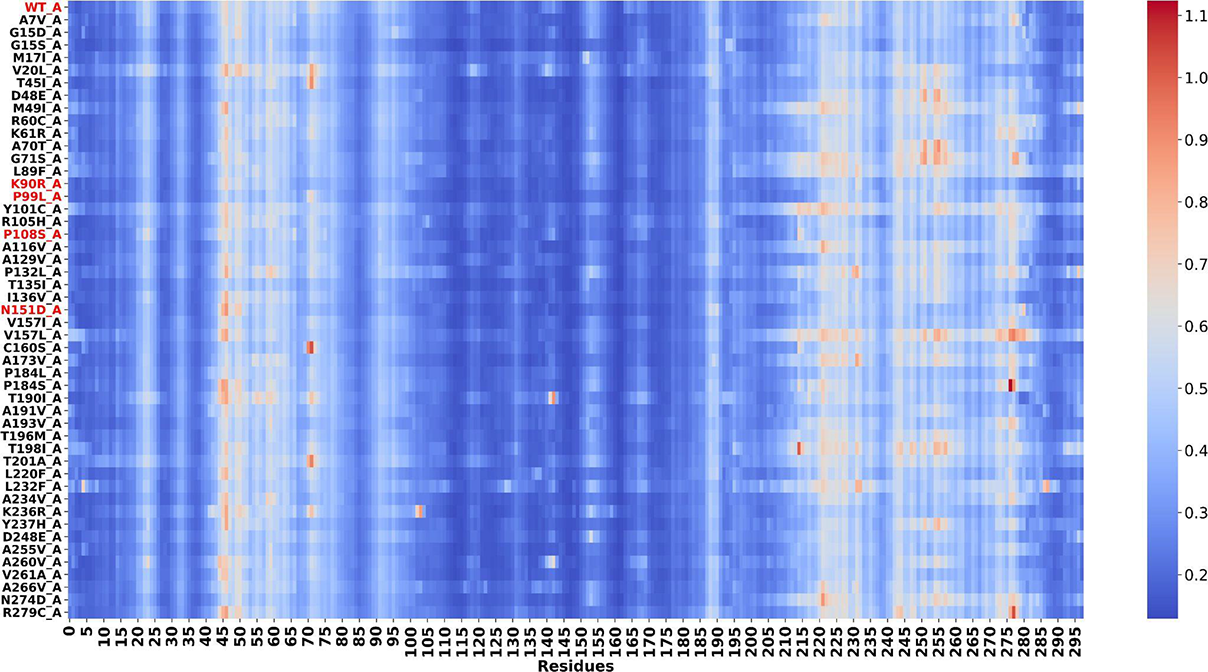
Heat map for Cα fluctuation of chain A of the WT and 48 mutants of SARS-CoV-2 M^pro^.

**Figure S3:**
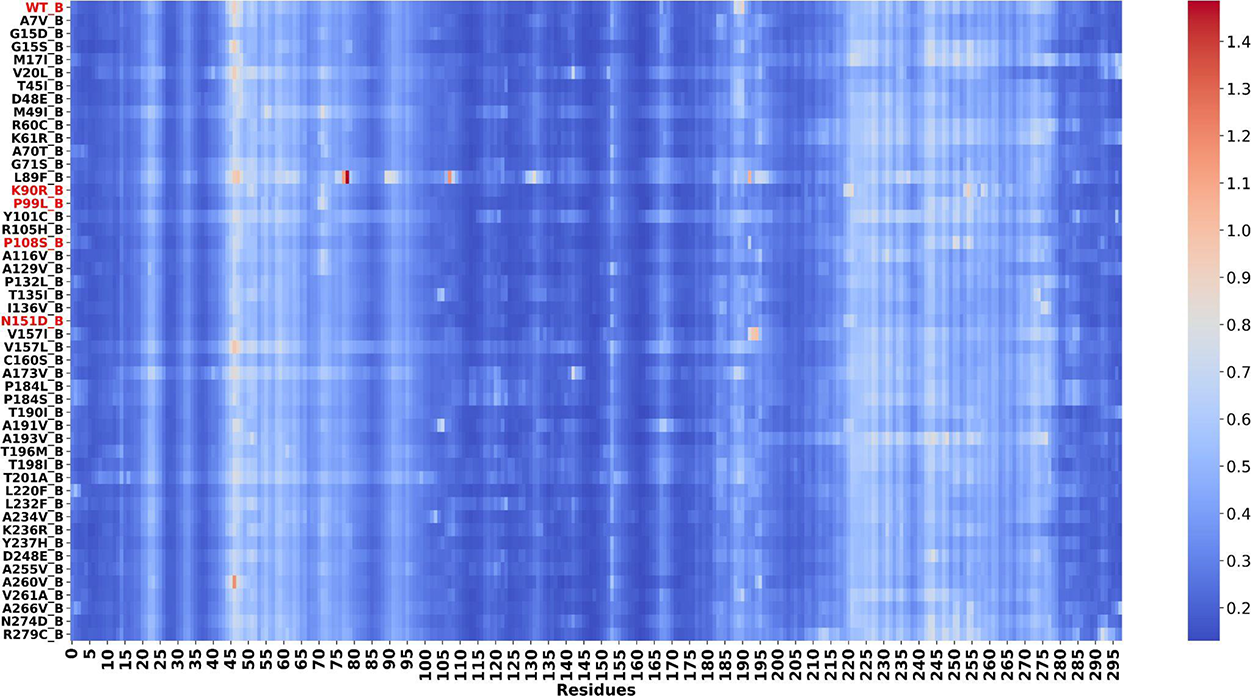
Heat map for Cα fluctuation of chain B of the WT and 48 mutants of SARS-CoV-2 M^pro^.

**Figure S4:**
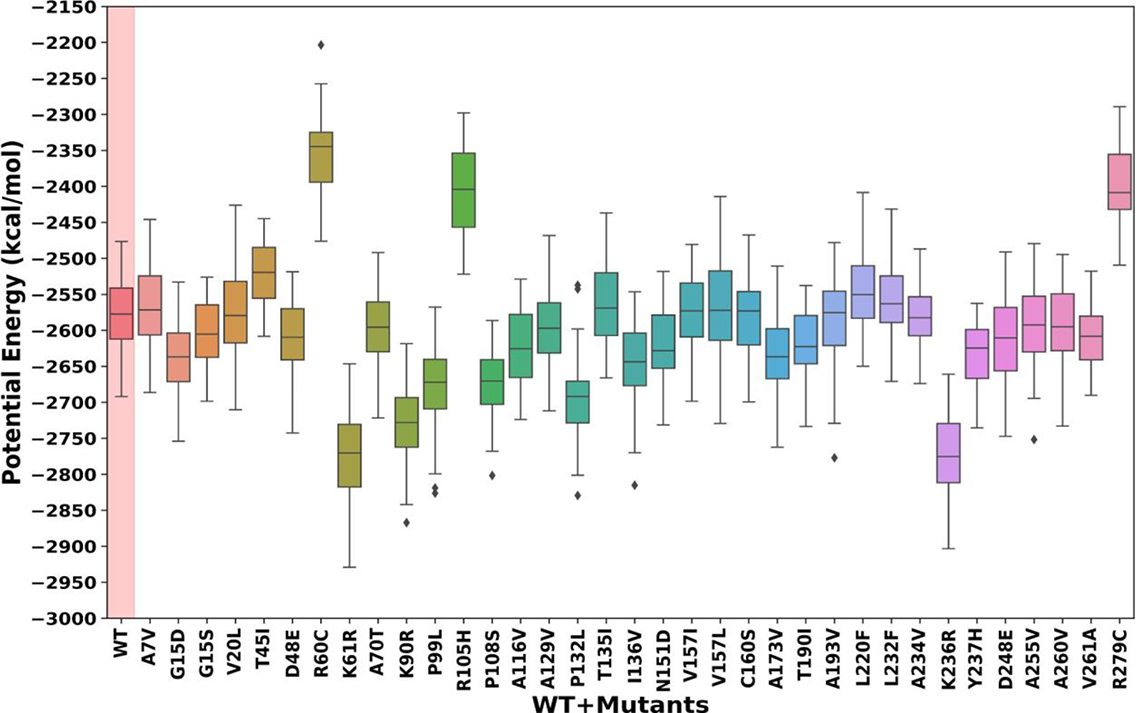
PotentialEnergy Boxplot for the structures generated by VMOD.The WT SARS-CoV-2 M^pro^ measure is highlighted by a red rectangle.

**Figure S5:**
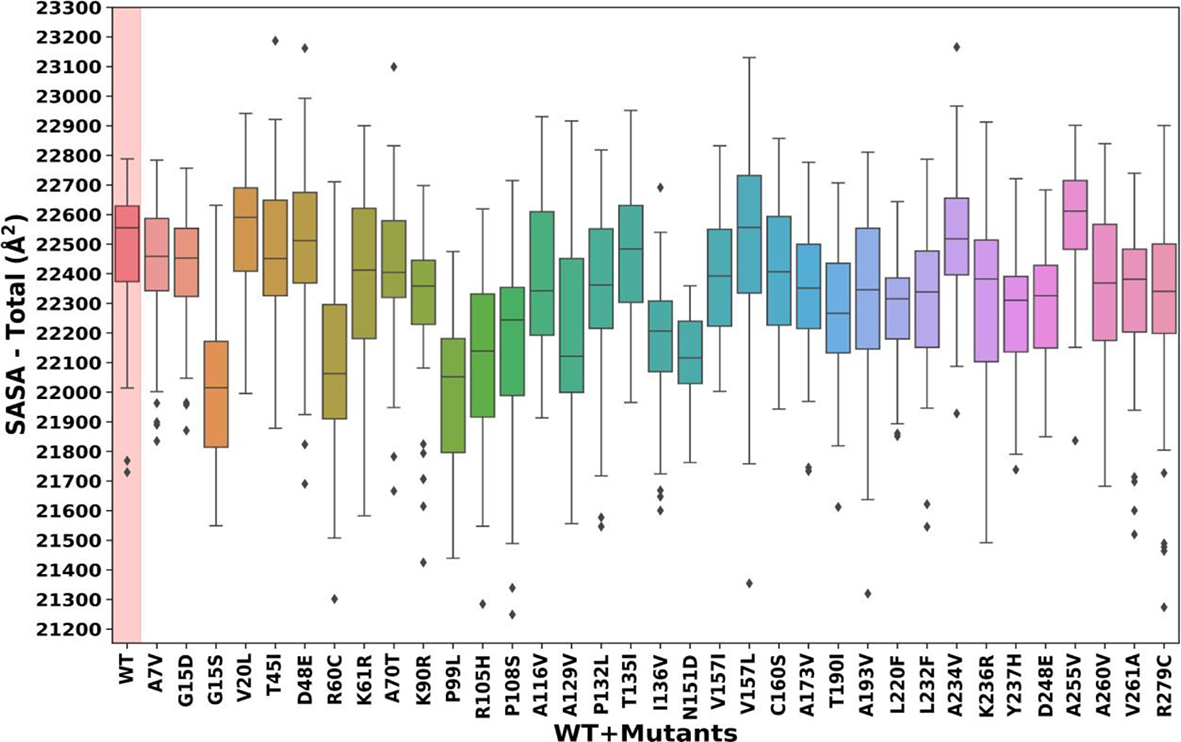
SASA Total Boxplot for the structures generated by VMOD. The WT SARS-CoV-2 M^pro^ measure is highlighted by a red rectangle.

**Figure S6:**
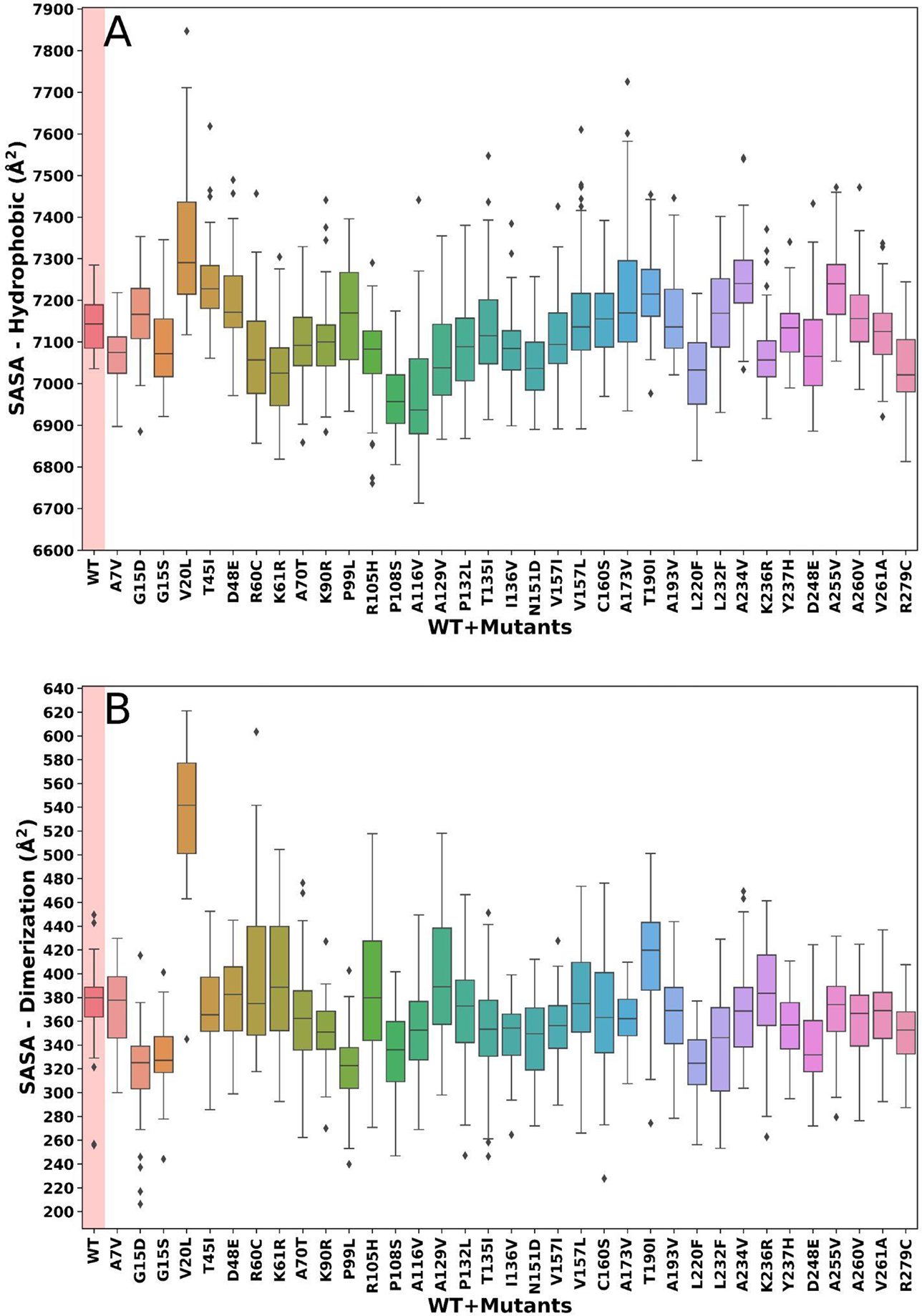
(A) Hydrophobic residues SASA and **(B)** Dimeric interface residues SASA Boxplot for the structures generated by VMOD. The residues involved in the dimer interface are listed in Table 1. The WT SARS-CoV-2 M^pro^ measure is highlighted by a red rectangle.

**Figure S7:**
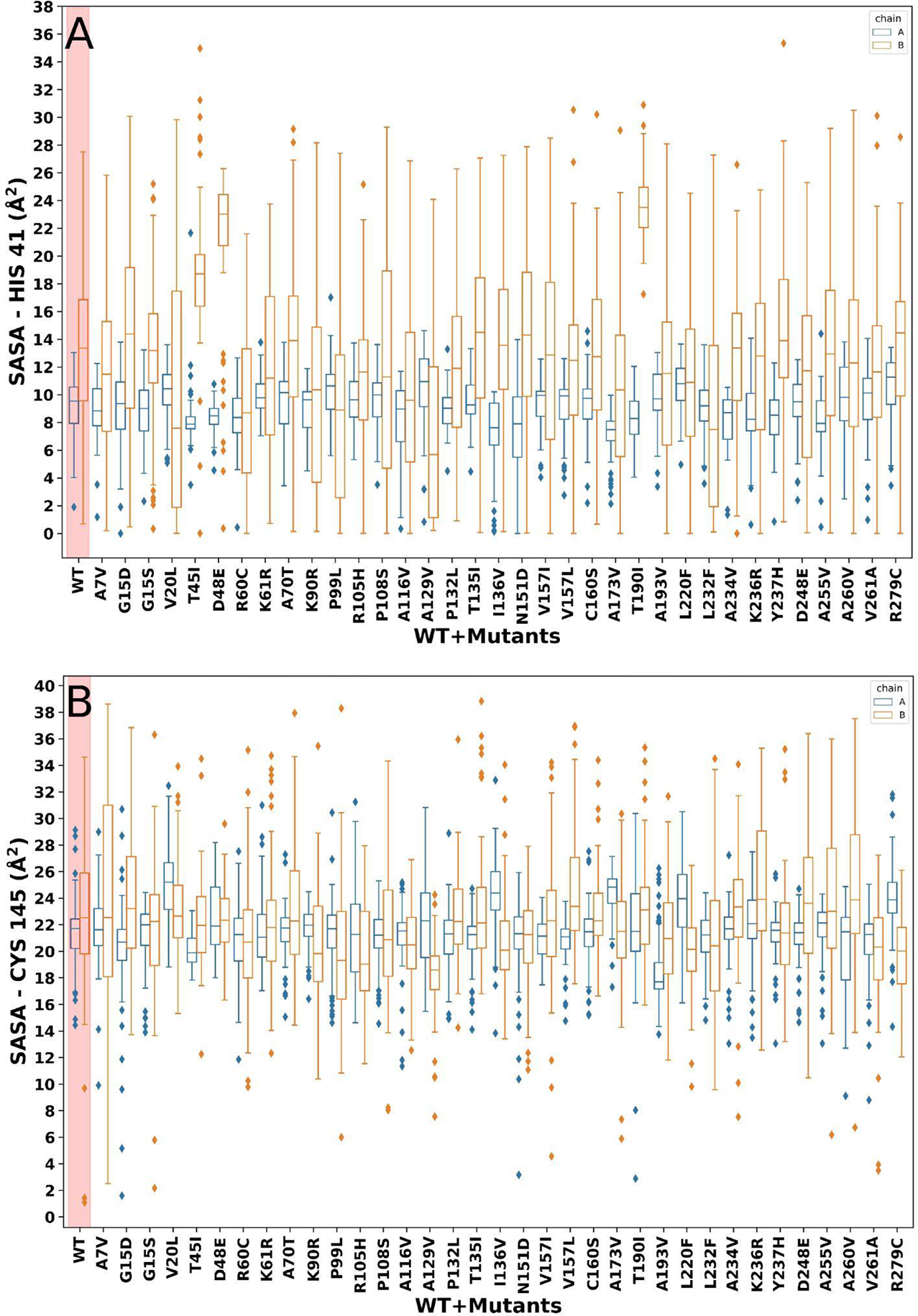
(A) SASA of His 41 and **(B)** SASA of Cys 145 catalytic dyad residues by chain. Chain A in blue and chain B in orange. The WT SARS-CoV-2 M^pro^ measures are highlighted by a red rectangle.

**Figure S8:**
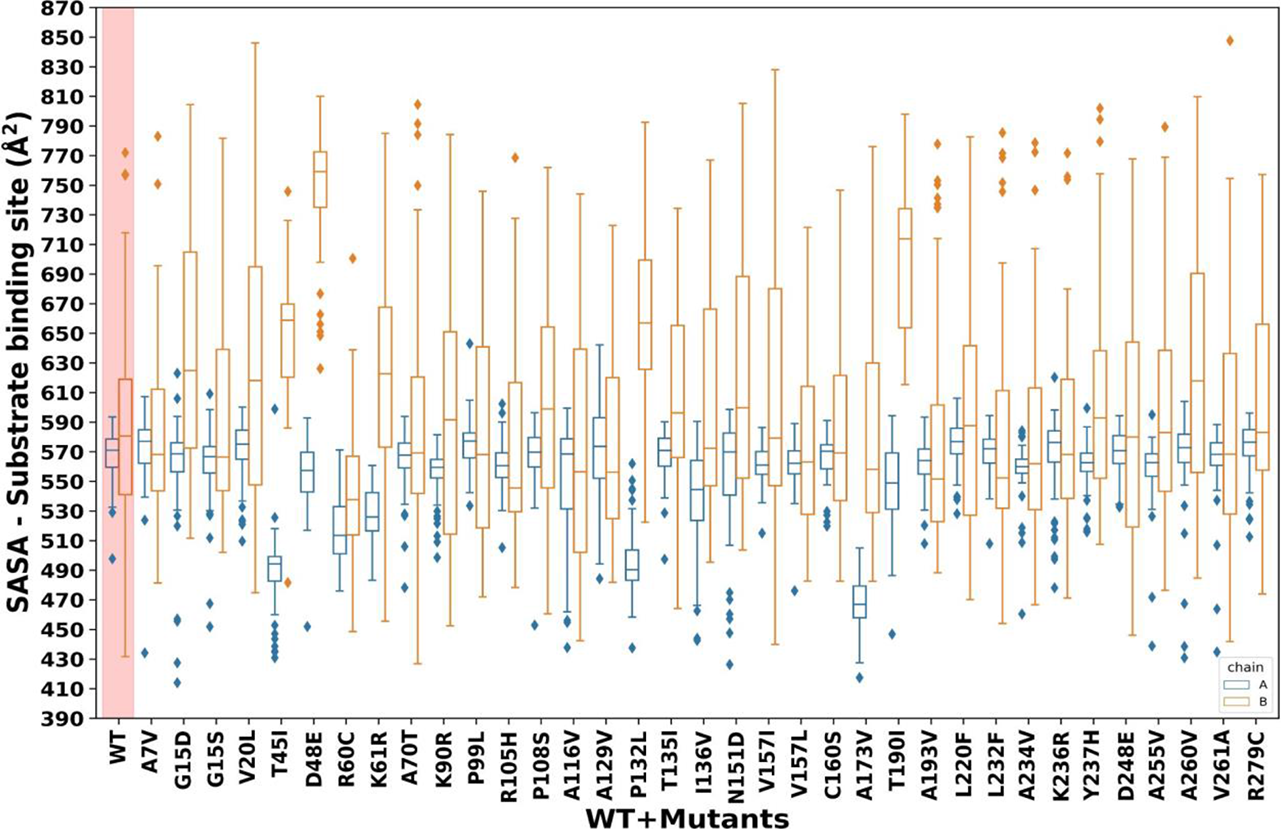
SASA of substrate binding sites presented by chain. Chain A in blue and chain B in orange. The WT SARS-CoV-2 M^pro^ measure is highlighted by a red rectangle.

**Figure S9:**
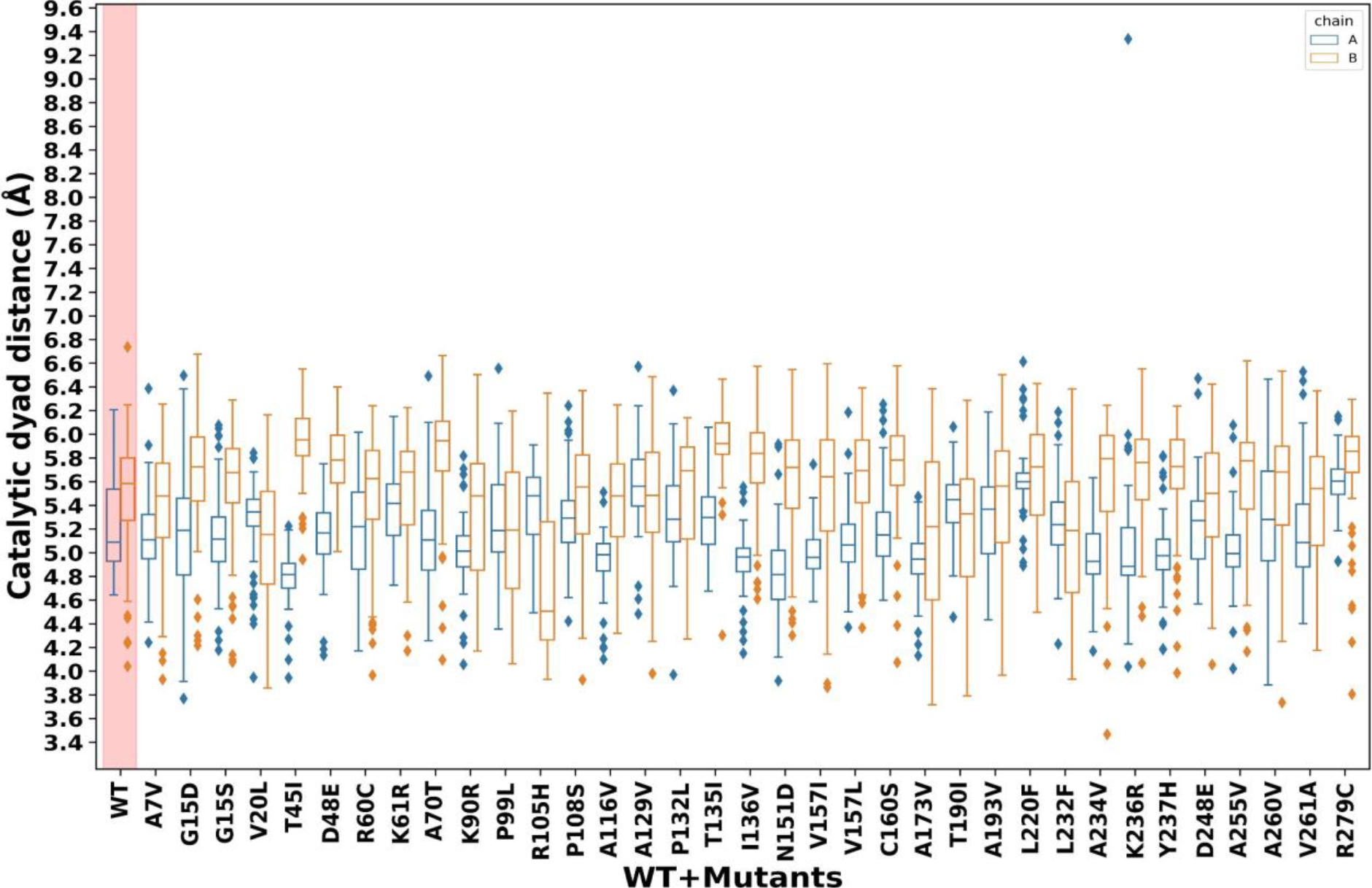
Catalytic Distance Boxplot by chain. Distance between the Cys 145 Sγ atom and His 41 Nε atom for both chains of the structures generated by VMOD. Chain A in blue and chain B in orange. The WT SARS-CoV-2 M^pro^ measures are highlighted by a red rectangle.

**Figure S10:**
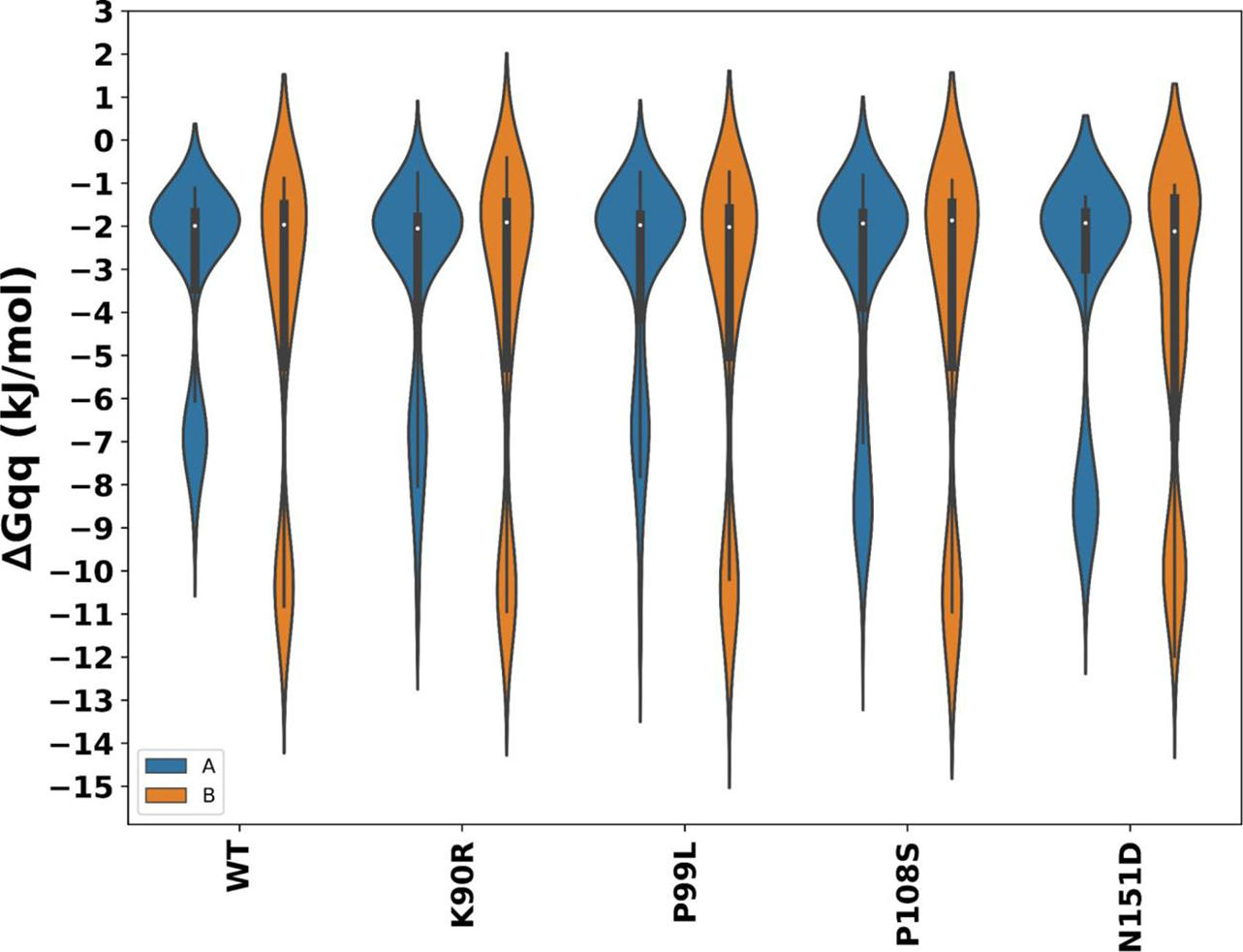
TKSA-MC ΔGqq (kJ/mol) of the interface residues Arg4, Glu14, Glu 290 and Arg 298 for WT and four mutants SARS-CoV-2 M^pro^.

**Figure S11:**
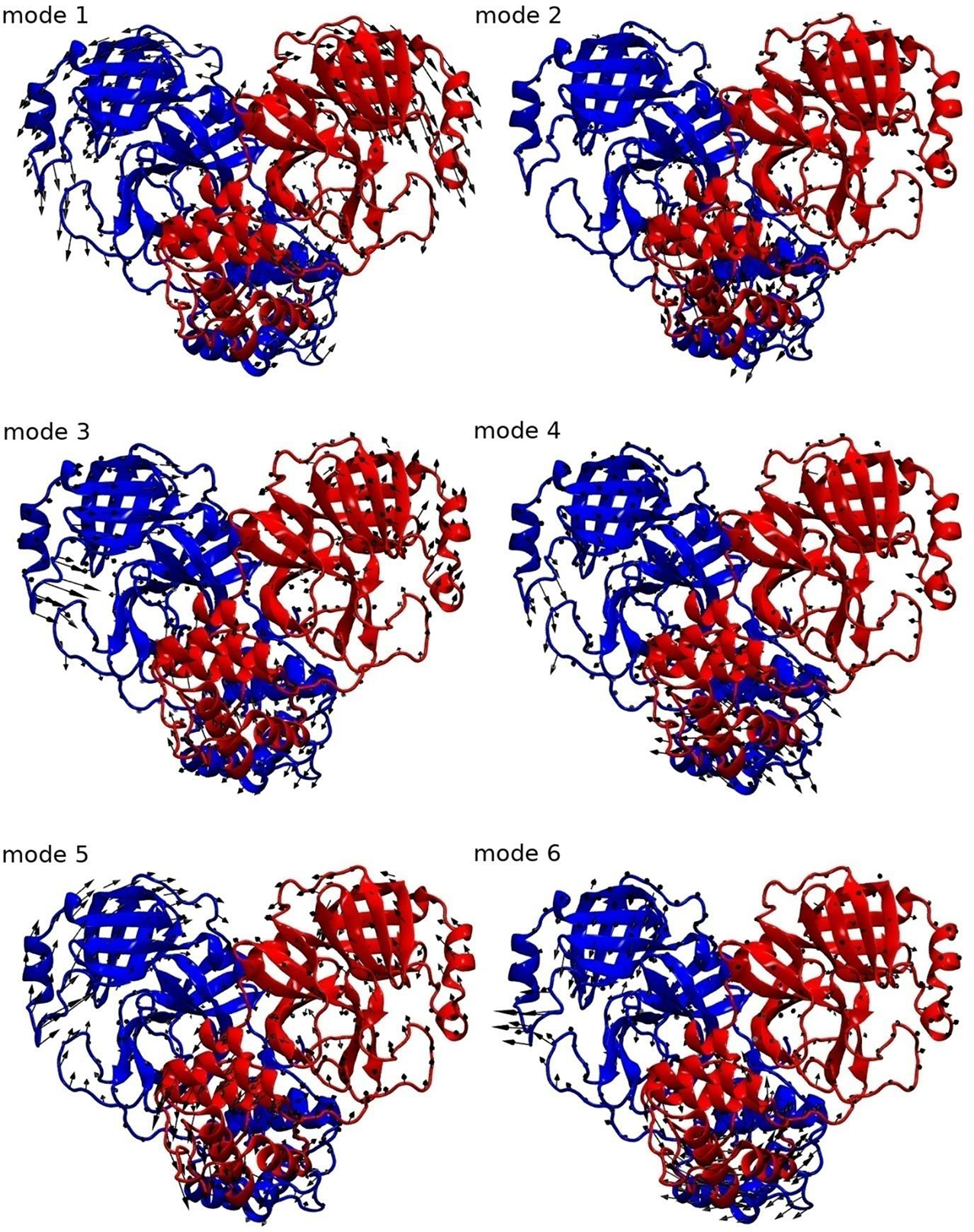
Representation of WT SARS-CoV-2 M^pro^ normal modes. The directions and the amplitude of the movements are represented by arrows. Figure generated in VMD [1].

**Table S1:** M^pro^ catalytic dyad distance (Å) for some dimeric apo structures of SARS-CoV-1 and SARS-CoV-2 available in PDB. Distance measured in PyMOL.

| **PDB ID** | **Virus** | **chain A** | **chain B** |
| --- | --- | --- | --- |
| 6WTM [2] | SARS-CoV-2 | 3.6 | 3.7 |
| 7ALI [3] | SARS-CoV-2 | 3.7 | 3.7 |
| 7C2Y [4] | SARS-CoV-2 | 3.7 | 3.9 |
| 7C2Q [5] | SARS-CoV-2 | 3.9 | 4.0 |
| 7LKD [6] | SARS-CoV-2 | 3.5 | 3.7 |
| 7CAM [7] | SARS-CoV-2 | 3.6 | 3.8 |
| 7MBG [8] | SARS-CoV-2 | 3.7 | 3.8 |
| 7CWC [9] | SARS-CoV-2 | 4.1 | 4.3 |
| 7KFI [8] | SARS-CoV-2 | 3.8 | 3.8 |
| 7BB2 [10] | SARS-CoV-2 | 3.8 | 3.8 |
| 6XHU [11] | SARS-CoV-2 | 3.8 | 3.8 |
| 3VB3 [12] | SARS-CoV-1 | 3.9 | 3.8 |
| 2DUC [13] | SARS-CoV-1 | 3.6 | 4.1 |
| 1UJ1 [14] | SARS-CoV-1 | 3.9 | 3.8 |
| 1UK3 [14] | SARS-CoV-1 | 3.9 | 3.8 |
| 1UK2 [14] | SARS-CoV-1 | 3.8 | 3.4 |
| 1Q2W [15] | SARS-CoV-1 | 3.8 | 3.9 |
| 2GT7 [16] | SARS-CoV-1 | 3.5 | 3.7 |

**Table S2:** Results from Mantel teste showing the modes of WT SARS-CoV-2 M^pro^ and their matching with the modes of the four mutants.

|  | **Mutants** |  |  |  |
| --- | --- | --- | --- | --- |
| **Wild-Type** | **K90R** | **P99L** | **P108S** | **N151D** |
| **mode 1** | no matching | no matching | no matching | no matching |
| **mode 2** | no matching | no matching | no matching | no matching |
| **mode 3** | no matching | mode 4 | no matching | mode 2 |
| **mode 4** | modes 2 and 4 | mode 4 | mode 4 | mode 5 |
| **mode 5** | mode 3 | mode 5 | no matching | no matching |
| **mode 6** | no matching | no matching | no matching | no matching |

